# Responses of global waterbird populations to climate change vary with latitude

**DOI:** 10.1101/784900

**Authors:** Tatsuya Amano, Tamás Székely, Hannah S. Wauchope, Brody Sandel, Szabolcs Nagy, Taej Mundkur, Tom Langendoen, Daniel Blanco, Nicole L. Michel, William J. Sutherland

## Abstract

While climate change continues to present a major threat to global biodiversity and ecosystems, most research on climate change impacts do not have the resolution to detect changes in species abundance and are often limited to temperate ecosystems. This limits our understanding of global responses in species abundance—a determinant of ecosystem function and services—to climate change including in the highly-biodiverse tropics. We address this knowledge gap by quantifying abundance responses to climate change in waterbirds, an indicator taxon of wetland biodiversity, at 6,822 sites between −55° and 64°. Using 1,303,651 count records since 1990 of 390 species, we show that with temperature increase, the abundance of species and populations decreased at lower latitudes, particularly in the tropics, but increased at higher latitudes. These contrasting responses to temperature increase according to latitude indicate potential global-scale poleward shifts of species abundance under climate change, providing empirical support for predictions by earlier studies. The negative responses to temperature increase in tropical species and populations are of conservation concern, as they are often also threatened by other anthropogenic factors. Our results suggest that existing biases in studies towards temperate regions could underestimate the impact of climate change on waterbirds and other species.

## Introduction

Climate change continues to pose various serious threats to biodiversity, and there is an urgent need to understand how species respond to changing climates globally. A wide range of species have already shown responses to climate change, such as changes in geographical range^1^, phenology^2^ and abundance^3^. However, the rate and direction of these responses vary greatly among species and locations^1,2,4^. As climate-driven changes in biodiversity are expected to affect ecosystem functioning, human well-being, and the dynamics of climate change itself^5^, understanding how species’ responses to climate change may vary globally could provide crucial evidence for a more effective allocation of limited resources on a priority basis for the conservation of species and ecosystems threatened the most by climate change, and for assessing how climate-driven changes in biodiversity may affect human societies.

Existing gaps in the geographical coverage of available evidence seriously limit our understanding of species’ responses to climate change and its variations across the globe^6^. Earlier global reviews of species’ responses to climate change have rarely incorporated species and studies in the tropics, due to the lack of relevant information^7^. Such geographical biases are even more prominent in studies investigating responses in species abundance^8,9^, which is a major determinant of species extinction risk^10^, ecosystem function and services^11^. Research on abundance responses to climate change to date has largely been conducted in Europe, North America and the Arctic^3,12-14^, with a recent global study showing a link between climate warming and abundance declines in birds and mammals^8^. As a result, although tropical species are predicted to be more vulnerable to increasing temperature^15^, there is still little empirical evidence on how responses in species abundance to climate change vary among and within species at the global scale.

Here we address this challenge by modelling global time-series data of waterbird species to estimate their abundance responses to changes in temperature and precipitation. The global dataset of waterbird abundance changes used is based on long-term surveys in over 100 countries and covers regions for which there is little information on climate change impacts, such as the tropics^16^. Waterbirds can also serve as an indicator taxon for assessing the status of biodiversity in wetland ecosystems, which have been lost at higher rates than other ecosystems, despite their high levels of biodiversity and productivity as well as the crucial ecosystem functions and services delivered^17^.

Using 1,303,651 count records collected since 1990 on 390 waterbird species at 6,822 sites between −55° and 64° (Figure S1) we first estimated, for each species at each site, (i) the rate of abundance changes with increasing temperature and precipitation as regression coefficients (responses to temperature and precipitation increases), and (ii) the proportion of abundance changes that can be explained independently by temperature and precipitation changes (measured as *R*^*2*^), estimated with hierarchical partitioning^18^ (the importance of temperature and precipitation). We then tested multiple hypotheses that are rarely explored at the global scale (Supplementary Tables S1 and S2), to examine among- and within-species variations in responses to temperature and precipitation increases as well as the importance of temperature and precipitation across latitudes.

## Results

Applying the Gompertz model of population growth to the global waterbird dataset enabled us to estimate abundance responses to the changes in temperature and precipitation at 1° × 1° grid cells across latitudes, including the tropics, for a wide range of waterbird groups. Of the 390 species analysed, 144 species (36.9%) had at least one estimate in the tropics and 129 species (33.1%) had their absolute latitudinal range mid-points in the tropics (defined as tropical species; Figure 1).

**Fig. 1.**
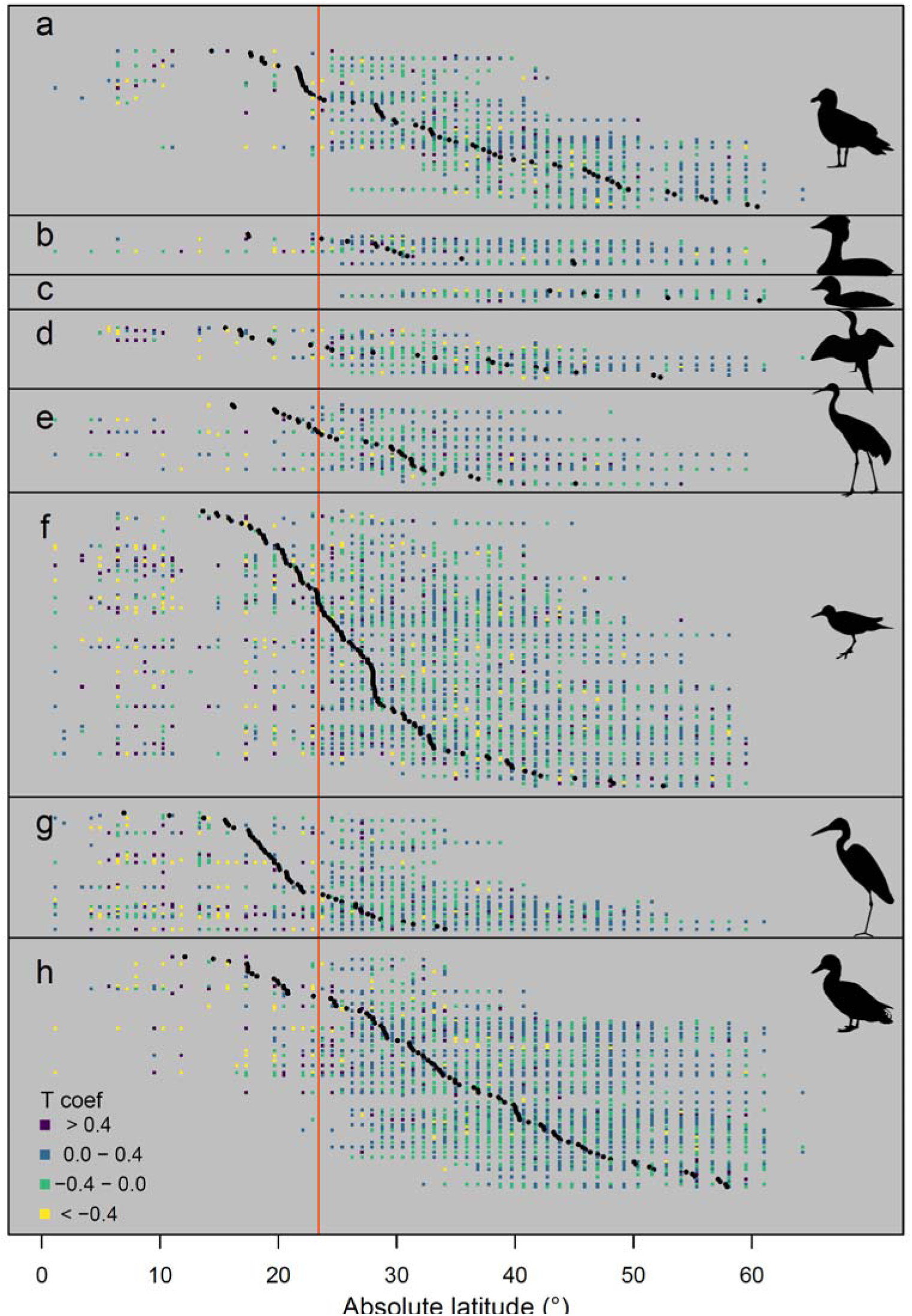
Latitudinal distribution of abundance responses to changes in temperature (the rate of abundance changes with increasing temperature) for each species. Each horizontal row of squares shows the absolute latitudes of 1° × 1° grid cells with estimates for each of the 390 species in (a) coursers, gulls, terns and auks, (b) grebes and flamingos, (c) loons and petrels, (d) pelicans, boobies and cormorants, (e) rails and cranes, (f) shorebirds, (g) storks, ibises and herons, and (h) waterfowl (see Methods for the definition of each species group). Black circles indicate the latitudinal range mid-point (i.e., median absolute latitude of geographical range) of each species. The area on the left of the red vertical line (absolute latitude < 23.4°) represents the tropical region.

Many species showed considerable spatial variation in abundance responses to temperature increases within their geographical ranges, with particularly negative responses in the tropics (Figure 1), although the importance of temperature in explaining abundance changes tended to be low across the ranges, with an overall median *R*^*2*^ of 0.057 (Supplementary Data S1 and S2). In contrast, for most species there was no clear geographical pattern in abundance responses to precipitation increases, and precipitation was found to have a low importance in explaining abundance changes (the overall median *R*^*2*^ = 0.051; Supplementary Data S1 and S2). These geographical patterns were also evident in the distribution of abundance responses averaged across all species observed within each grid cell; species generally showed more negative responses to temperature increases at lower latitudes, such as in South and Southeast Asia (Figure 2).

**Fig. 2.**
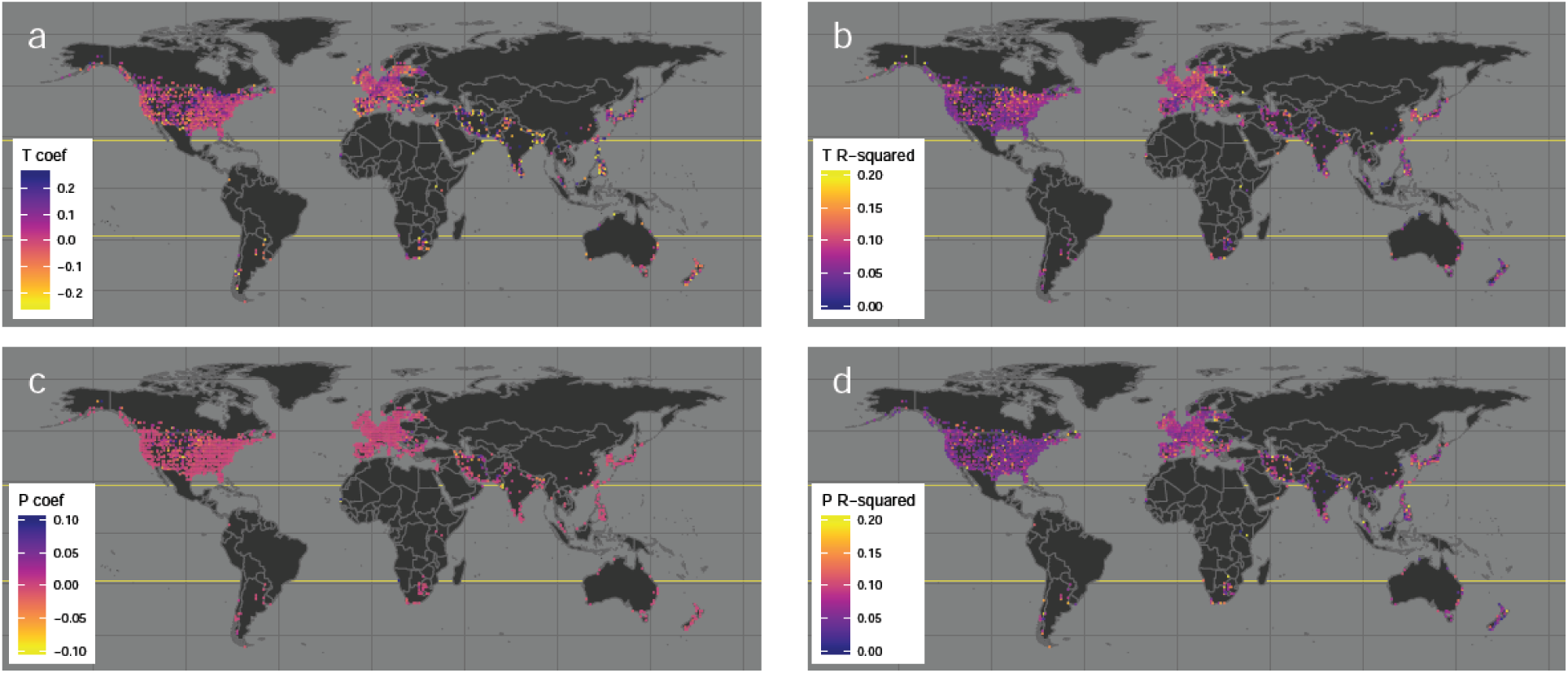
Mean abundance responses across 390 waterbird species to changes in temperature and precipitation in each 1° × 1° grid cell. The rate of abundance changes with increasing temperature (a), the independent capacity of temperature in explaining abundance changes (b), the rate of abundance changes with increasing precipitation (c) and the independent capacity of precipitation in explaining abundance changes (d). The region between the yellow solid lines is the tropics.

For 213 species with estimates at ten or more grid cells, we then tested hypotheses on how responses to temperature and precipitation increases (the rate of abundance changes with increasing temperature/precipitation) and the importance of temperature and precipitation (the proportion of abundance changes that can be explained by temperature/precipitation changes) vary among species (among each species’ estimates at latitudinal range mid-points; species-level responses) and within species (among grid cells within each species; population-level responses) along latitudes. When compared among species, abundance responses to temperature increases were more negative in species at lower latitudes, with 69% of the tropical species showing negative responses to temperature increases (Figure 3a, S2a). When compared within species, although 198 (93%) out of the 213 species showed more negative responses to temperature increases at lower latitudes, this within-species latitudinal pattern was significant only in eight of the 198 species (Figure 3b, Supplementary Data S3). The importance of temperature in explaining abundance changes also increased with latitude among species (Figure 3c, S2b) and within species for all 55 species with a significant within-species latitudinal pattern (Figure 3d, Supplementary Data S3). Migratory species, larger-sized species and species with a wider latitudinal range showed a higher importance of temperature in explaining abundance changes (Figure S2b).

**Fig. 3.**
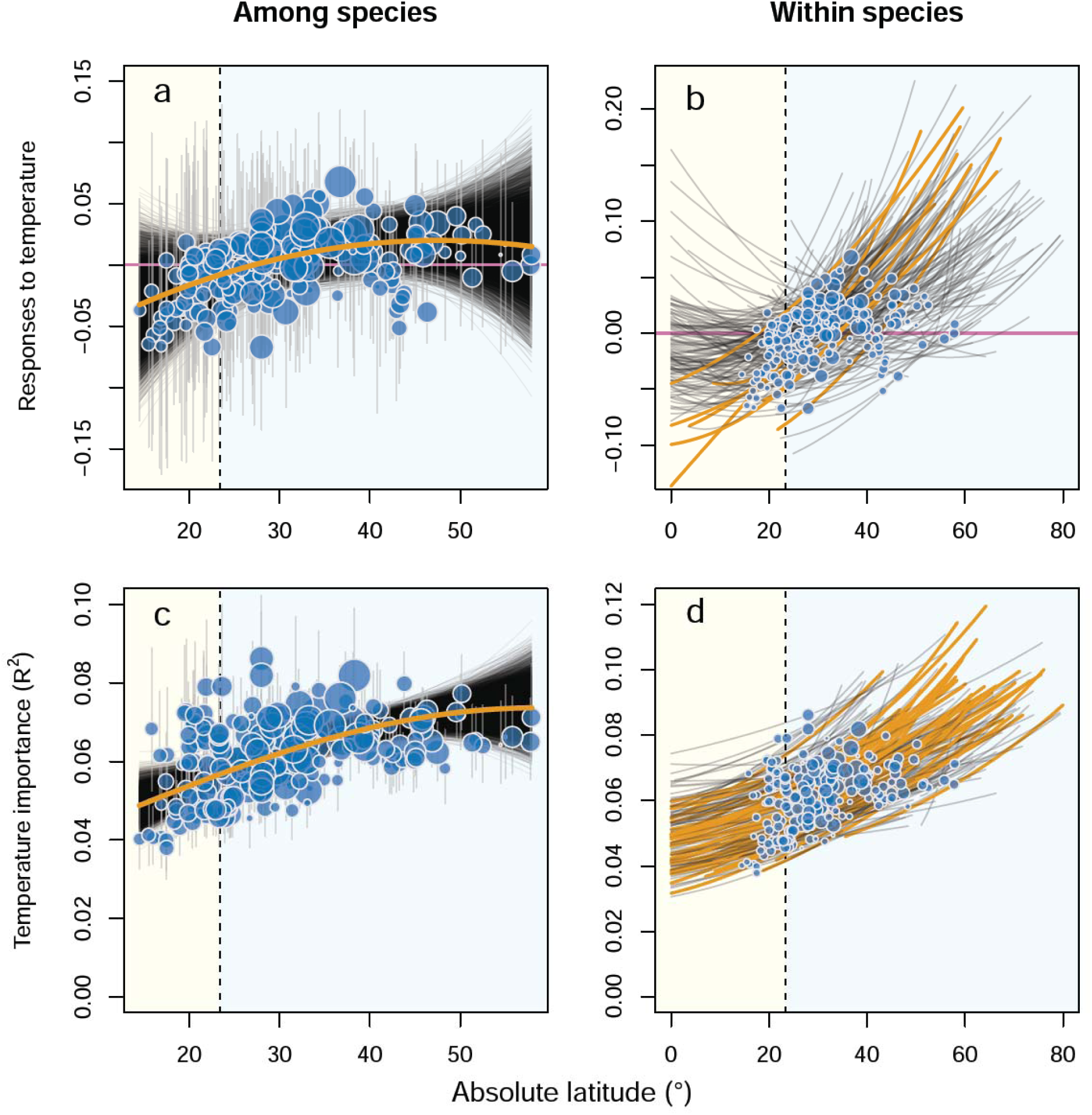
Latitudinal patterns in waterbird abundance responses to temperature increases. The rate of abundance changes with increasing temperature at each species’ range mid-points (a) and within each species (b), the independent capacity of temperature in explaining abundance changes at each species’ range mid-points (c) and within each species (d). In (a) and (c) orange lines represent the among-species latitudinal patterns based on posterior median coefficients with black lines showing patterns based on all posterior samples. Blue circles indicate the estimated responses at each species’ range mid-points (also shown in (b) and (d)) with grey vertical lines showing 95% credible intervals; circle size, the absolute latitudinal range size. In (b) and (d) regression lines show the estimated within-species latitudinal patterns for each species based on posterior median coefficients; orange lines represent significant patterns. The yellow area (absolute latitude < 23.4°) represents the tropical region.

In contrast, neither abundance responses to precipitation increases nor the importance of precipitation in explaining abundance changes showed significant latitudinal patterns among species, although, for some species in the tropics, precipitation was found to have a relatively high importance in explaining abundance changes (Figure 4a, c, S3 and Supplementary Data S3). Precipitation was shown to have a higher importance in explaining abundance changes in species with a wider latitudinal range (Figure S3b). When compared within species, three species showed a significant, one species showed a decrease and another species showed a hump-shaped curve in abundance responses to precipitation increases along latitudes (Figure 4b, Supplementary Data S3). The importance of precipitation in explaining abundance changes showed a significant within-species latitudinal pattern for just one species (Figure 4d, Supplementary Data S3).

**Fig. 4.**
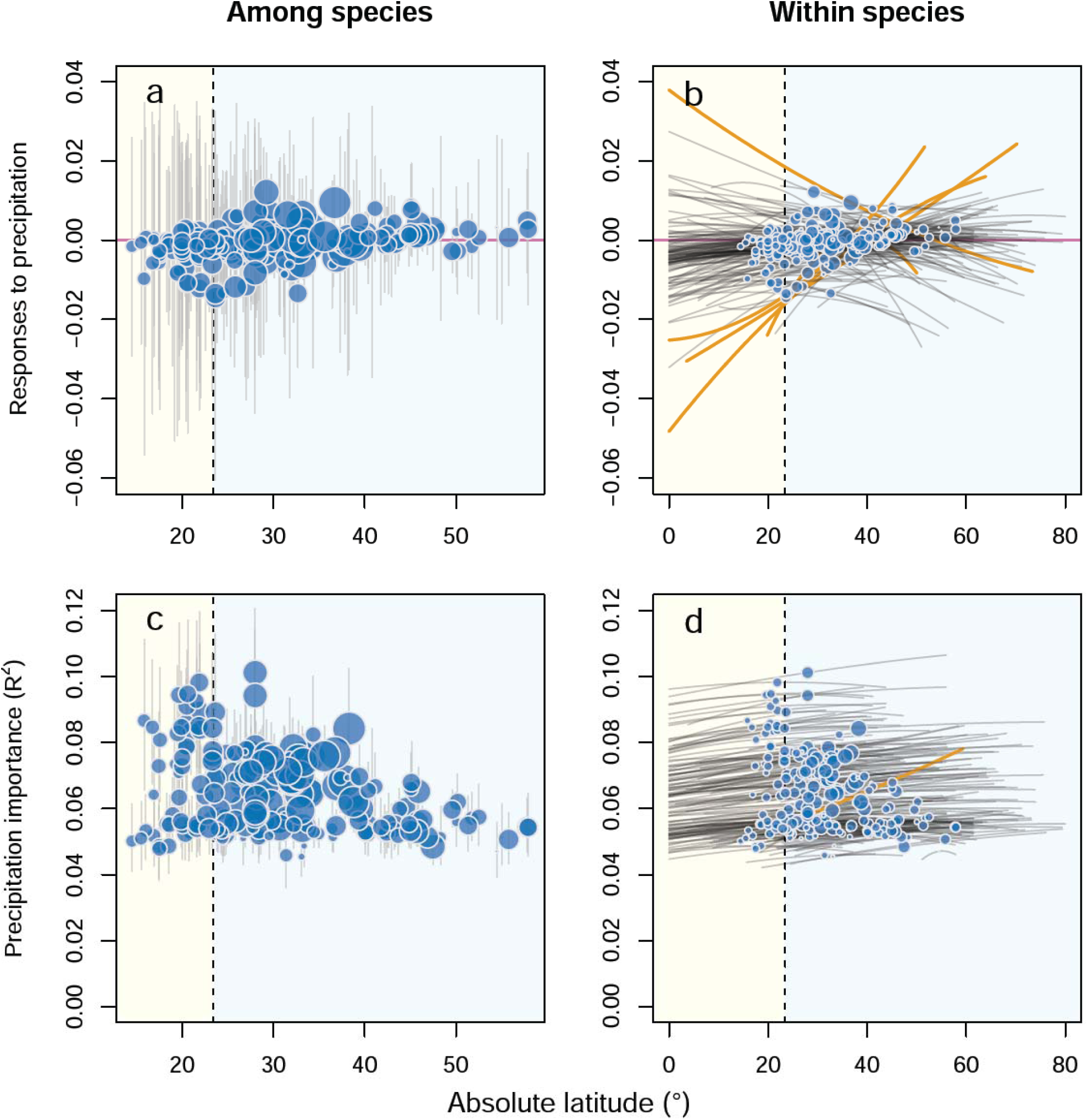
Latitudinal patterns in waterbird abundance responses to precipitation increases. The rate of abundance changes with increasing precipitation for each species’ range mid-points (a) and within each species (b), the independent capacity of precipitation in explaining abundance changes at each species’ range mid-points (c) and within each species (d). In (a) and (c) orange lines represent the among-species latitudinal patterns based on posterior median coefficients with black lines showing patterns based on all posterior samples. Blue circles indicate the estimated responses at each species’ range mid-points (also shown in (b) and (d)) with grey vertical lines showing 95% credible intervals; circle size, the absolute latitudinal range size. In (b) and (d) regression lines show the estimated within-species latitudinal patterns for each species based on posterior median coefficients; orange lines represent significant patterns. The yellow area (absolute latitude < 23.4°) represents the tropical region.

## Discussion

Our results demonstrate the responses in waterbird abundance to temperature increases differ between tropical and non-tropical regions. At both species and population levels, waterbird abundance generally decreased in the tropics, but increased at higher latitudes, with increasing temperature. This supports our predictions on among- and within-species patterns (Supplementary Table S1). Species in the tropics tend to live closer to their upper temperature limits^15^, have a narrower temperature niche^19^ and change their temperature niche at a slower rate^20^, all of which indicate that tropical species are more vulnerable to increasing temperatures at the species level. Climate-related extinctions of local populations, typically at the warmer edge of the species’ geographical range, are also more frequent in the tropics, causing poleward range shifts in many species^21^. While such species-level and population-level responses to climate change have often been investigated separately to date, our results provide novel empirical evidence that impacts of temperature increases on tropical ecosystems can be characterised by species-wide declines in tropical species as well as population-level responses in wider-ranging species.

Nevertheless, we also found that temperature generally explains only a small proportion of yearly abundance changes in waterbirds, especially in tropical species and at the low-latitude range margin of species, possibly for four reasons. First, the effect of temperature changes on waterbird abundance can be indirect especially at lower latitudes. Although warmer weather conditions can directly increase the survival of waterbirds at higher latitudes^22^, indirect biotic processes (e.g., changes in food availability), rather than direct abiotic processes (e.g., heat stress), are reported to be more important mechanisms for climate-driven abundance changes, especially for higher-level consumers like birds^23,24^. For example, increases in already-high temperatures at lower latitudes can cause wetlands to dry, reducing the availability of habitats and food resources for waterbirds^22^. Such an indirect effect of temperature increases could have obscured the relationship between changes in temperature and abundance, especially in the tropics and at the low-latitude range margins. Second, many of the waterbirds analysed here are migratory species, which generally have a higher dispersal ability^25^ and track climate niches to a greater extent than resident species^26^, and thus can be more responsive to changes in local temperature; this was supported by the positive effect of migratory status on the importance of temperature (Figure S2b). In this study more non-tropical species tended to be migratory compared to tropical species (151 (96%) of 158 non-tropical species and 12 (78%) of 55 tropical species were migratory), which may explain the higher importance of local temperature in explaining the abundance of non-tropical species. Third, observation errors can cause a lower explanatory power of variables. We may expect larger errors in the tropics, compared to temperate regions, where waterbird surveys have a longer history and thus surveyors might be better trained. Finally, other important threats, such as habitat loss and hunting, affect bird abundance, independently from, or synergistically with, climate change^27^. By testing the effect of temperature and precipitation changes on yearly abundance changes while estimating long-term growth rates, our modelling approach controlled for the consistent impacts of such threats on long-term trends in abundance (see *Statistical Analyses* for more detail). Nevertheless, those threats can also cause yearly abundance changes and their impacts are likely to be more severe at lower latitudes^16^, potentially causing temperature to have lower importance at lower latitudes.

Contrary to our hypotheses, there was no clear latitudinal pattern in abundance responses to precipitation changes, either among or within species. In general water availability, compared to ambient temperature, has been shown to be a more important driver of species richness and population size at lower latitudes^4,28^. Supporting this, our results showed that precipitation was more important in explaining the abundance of some tropical species compared to most species in higher latitudes, although the overall among-species pattern across latitudes was not significant (Figure 4c). This may be explained by the following two reasons. First, precipitation changes can affect waterbird species at the river basin scale (often the scale of 500 to 1,000 km) through effects on water flow into their wetland habitats^29,30^. Therefore, our analysis at the resolution of 1° grid cells (equivalent to a grain size of 96.49 km) may not have been able to detect such a broad-scale impact of precipitation changes. Second, waterbird responses to precipitation changes can vary greatly among species, as we recognised when developing our hypotheses (Supplementary Table S1). While increased rainfall generally leads to more favourable habitat conditions for waterbirds in dry regions^22,31^, elevated water levels associated with increased rainfall can cause the loss of shallow-water habitats, often followed by abundance decreases in certain groups, such as shorebirds^32^. Such mixed responses to precipitation changes among species^33^ may have resulted in the lack of clear latitudinal patterns, particularly among species, in this study.

Our results point to three major implications on the impact of climate change on global biodiversity. First, despite the relatively low importance of temperature in explaining waterbird abundance changes, all other things being equal, local temperature increases under ongoing climate change are likely to pose a more negative impact on species and populations in the tropics. This provides important evidence for the climate-driven degradation of tropical ecosystems, which has recently been under debate^34,35^. Although climate change is not the only threat to waterbird species, impacts of other major threats, such as loss and degradation of wetlands and excessive hunting pressure, seem to be more severe in the tropics too^16^, indicating that tropical species and populations suffer from multiple anthropogenic threats. Second, the revealed negative impact of temperature increases in the tropics suggests that existing severe biases in scientific studies and data towards temperate regions could underestimate the impact of climate change on species populations at the global scale. Highlighting the negative impact of climate change on tropical waterbirds should serve to inspire further studies on other taxa in the tropics. Finally, our finding of contrasting abundance responses to temperature increases according to latitude highlights the possibility of global-scale poleward shifts in abundance across species, and associated ecosystem functions and services. As such shifts can have serious consequences not only for biodiversity but also for human well-being, assessing latitudinal patterns in biodiversity responses to climate change at the population, species and community levels warrants further research attention.

## Methods

### Data

#### Waterbird count data

Data used in this study comprised site-specific annual counts based on the International Waterbird Census (IWC) coordinated by Wetlands International and the Christmas Bird Count (CBC) by the National Audubon Society in the USA, and were compiled in our earlier study^16^. Note that counts based on these surveys should be described as relative abundance, as we could not account for imperfect detections in this study. However, we have referred to them as abundance throughout the manuscript for simplicity. Nevertheless, these count records should still be comparable among years (see section *Model for estimating abundance responses* for more detail).

The IWC, launched in 1967, is a scheme for monitoring waterbird numbers, covering more than 25,000 sites in over 100 countries with more than 15,000 observers. The coordination of the IWC is divided into four regional schemes corresponding to the major migratory flyways of the world: the African-Eurasian Waterbird Census (AEWC), Asian Waterbird Census (AWC), Caribbean Waterbird Census (CWC) and Neotropical Waterbird Census (NWC). We did not use data from the CWC, as, having started in 2010, it only provides short-term data. The survey methodology is essentially the same across the four regional schemes. Population counts are typically carried out once every year in mid-January but may include counts between December to February. Additional counts are also conducted in other months, particularly in July in the Southern Hemisphere, but we only used the January and February counts for consistency. This means that our data from the Northern Hemisphere are for non-breeding populations while those in the Southern Hemisphere also include some breeding populations. In each country that is covered by the survey, national coordinators manage an inventory of wetland sites (hereafter, survey sites), including sites of international- or national-level recognition (e.g., Ramsar sites, Flyway Network Sites, Important Bird Areas, national parks etc.). Each survey site is generally defined by boundaries so that observers know precisely which areas are to be covered in the surveys. The observers consist of a wide variety of volunteers, but national coordinators usually train them using materials produced by Wetlands International to ensure the quality of count data. Survey sites (normally up to a few km^2^) are typically surveyed by about two observers for up to four hours, while larger sites can require a group of observers working over several days. Most surveys are conducted on foot, or from a vehicle, with boats involved in a few. The time of survey on any given day depends on the type of survey site: inland sites are normally surveyed during the morning or late afternoon, whereas coastal sites are surveyed over the high tide period (mangrove areas and nearby mudflats are, however, covered during low tides). Surveys cover waterbirds, defined as bird species that are ecologically dependent on wetlands^36^. Counts are usually made by scanning flocks of waterbirds with a telescope or binoculars and counting each species. Zero counts are not always recorded, and thus are inferred using a set of criteria (see below for more detail). Count records, together with associated information, are submitted to the national coordinators, who compile the submitted records, check their validity and submit those records to Wetlands International. See ^36,37^ for more details on survey methodology.

As the IWC does not cover North America, we also used data based on the CBC, which has been conducted annually since 1900, and now includes over 2,400 count circles (defined as survey sites in this study) and involves more than 70,000 observers each year^38^. Each CBC consists of a tally of all bird species detected within 24.1 km in diameter, on a single day between 14th December and 5th January. The majority of circles (and most historical data) are from the US and Canada. Observers join groups that survey subunits of the circle during the course of the day using a variety of transportation methods (mostly on foot, or in a car, but can include boats, skis, or snowmobiles). The number of observers and the duration of counts vary among circles and through time. The total number of survey hours per count has been recorded as a covariate to account for the variable duration of and participation in the count. We only used records on waterbird species in this paper.

We compiled data from each scheme by species, except for data based on the African-Eurasian Waterbird Census, where data had already been stored by flyway for each species^37^. As data based on the Neotropical Waterbird Census are only available for 1990 onward, we only used post-1990 data for other regions as well. The latest records were in 2013. For the IWC data, we generated zero counts using an established approach^37^, in which we started with a list of all species observed in each country and assumed a zero count of any species that were on the list but not recorded at a particular site on a particular day if the site was surveyed on that day, as shown by the presence of any other species’ record(s), and if no multi-species code related to the species (e.g., Anatinae spp. for species of the genus *Anas*) was recorded for the site-date combination. We projected all survey sites onto a Behrmann equal-area cylindrical projection and assigned them to grid cells with a grain size of 96.49 km, or approximately 1° at 30° N/S. We only used species that were observed at one or more survey sites for ten or more years since 1990, and this has resulted in 390 species being analysed in this study (see Supplementary Data S4 for the full list of species). Species groups used in Fig. 1 are based on the International Ornithological Congress World Bird List^39^: coursers, gulls, terns and auks (Alcidae, Glareolidae, Laridae and Stercorariidae), grebes and flamingos (Phoenicopteridae and Podicipedidae), loons and petrels (Gaviidae and Procellariidae), pelicans, boobies and cormorants (Anhingidae, Fregatidae, Pelecanidae, Phalacrocoracidae and Sulidae), rails and cranes (Aramidae, Gruidae and Rallidae), shorebirds (Burhinidae, Charadriidae, Haematopodidae, Jacanidae, Recurvirostridae, Rostratulidae and Scolopacidae), storks, ibises and herons (Ardeidae, Ciconiidae and Threskiornithidae), and waterfowl (Anatidae and Anhimidae).

#### Explanatory variables

To estimate responses in waterbird abundance to changes in temperature and precipitation, we used monthly mean temperature and precipitation total in the CRU TS v. 4.01 database ^40^, by assigning each site to the 0.5° climatic grid cell including the site. When testing among- and within-species latitudinal patterns in abundance responses, we also accounted for three species-level variables—latitudinal geographical range, migratory status and body size—that are expected to explain among-species variations in responses: data sources of those variables are shown in Supplementary Table S2.

### Statistical Analyses

#### Model for estimating abundance responses

We first estimated for each species at each survey site the rate of abundance changes with increasing temperature and precipitation as regression coefficients (defined as abundance responses to temperature or precipitation increases) by applying the Gompertz model of population growth to count records:

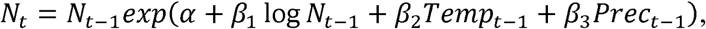

where *N*_*t*_, *Temp*_*t*_, *Prec*_*t*_ are the abundance of the species, the relevant mean Dec-Feb temperature and precipitation at the site in year *t*, respectively. *β*_*1-3*_ are regression coefficients and *α* is the intercept. By estimating *α* as the population growth rate, this model tests the effect of temperature and precipitation on yearly changes in abundance while controlling for long-term trends in abundance. This model structure helps to avoid detecting a spurious relationship between long-term trends in abundance caused by other threats (e.g., long-term declines by habitat loss) and those in temperature or precipitation (e.g., long-term warming temperatures). Taking logs and rearranging to express in terms of relative growth rate result in the following form:

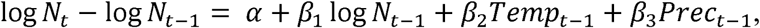

and we used this form to estimate regression coefficients with linear models in R 3.4.1^41^. As this model does not allow missing values, any missing values between the first and last survey years at each site for each species were replaced by linear interpolation using the package zoo^42^; the proportion of missing values (i.e., the effect of interpolation) was accounted for in the following analysis (see *Latitudinal analysis*). The estimated *β*_*2*_ and *β*_*3*_ represent site-level abundance responses to temperature and precipitation increases, respectively. Using the same model we also estimated the site-level independent capacity of temperature and precipitation changes in explaining abundance changes (defined as the importance of temperature and precipitation) with hierarchical partitioning^18^ (measured in our case as R^2^) using the package hier.part^43^.

As the model described above tests the effect of temperature and precipitation in the previous year (i.e., year *t-1*) on abundance in the survey year (year *t*), we also separately tested the immediate effect of temperature and precipitation in the same year (year *t*) as the abundance survey year. For this analysis we used mean temperature and precipitation in December (year *t-1*), January and February (year *t*) for the IWC sites, where surveys were conducted either in January or February, and mean Dec temperature and precipitation in year *t* for the CBC sites, where surveys were largely conducted in December. We compared AIC of the two models at each site for each species and used the coefficients in the model with a smaller AIC.

We assumed constant survey efforts over time for the IWC, because regular and standardized surveys with constant methods, efforts and timing are strongly encouraged in this scheme (see Supplementary Discussion in^16^ for more detail). However, survey efforts in the CBC are known to vary through time. Following a previously published analysis^44^ we thus accounted for the survey effort effect for the CBC data by using the total number of survey hours per count as the measure of survey efforts:

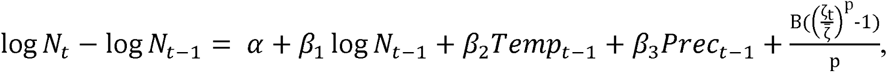

where *ζ*_*t*_ is the total number of survey hours per count and 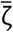 is the mean value of *ζ*_*t*_. The parameters B and p determine a range of relationships between effort and the number of birds counted^44^ and we used the values estimated for each species in our earlier study^16^ (see Supplementary Data S4).

We only used survey sites with ten or more records and five or more non-zero records since 1990 for at least one species, and this has resulted in 1,303,651 count records since 1990 on 390 species at 6,822 sites between −55° and 64° (Supplementary Figure S1) being analysed in this study. We aggregated the estimated site-level responses to temperature and precipitation increases as well as the importance of temperature and precipitation to 1°×1° grid cells by calculating the mean site-level estimates across all sites in each grid cell, weighted by the inverse of estimate variance at each site to account for uncertainties. The grid cell-level estimates (Supplementary Data S2) were then used in the latitudinal analysis described below and for the species-level maps (Supplementary Data S1). We also calculated community-level responses (Figure 2) by calculating the mean grid cell-level estimates across all species observed in each grid cell, weighted by the inverse of estimate variance in each species to account for uncertainties.

#### Latitudinal analysis

We used absolute latitudes to test latitudinal patterns described in Supplementary Table S1 for the following reason. Our data include species that are distributed only in either the northern or southern hemisphere (one-hemisphere species) as well as those that appear in both the hemispheres (two-hemisphere species). Some of our hypotheses (e.g., that for among-species patterns in abundance changes with increasing temperature, shown at the top of Supplementary Table S1) predict that one-hemisphere species would show a monotonic increase with raw latitudes while two-hemisphere species would show a U-shaped relationship along the raw latitudinal gradient with the lowest point at the equator; this makes analysing those species together in the hierarchical modelling framework described below a complicated process. With absolute latitudes, in contrast, one-hemisphere and two-hemisphere species are both expected to show a monotonic increase, making the parameter estimation much simpler.

To explain among- and within-species latitudinal variations in abundance responses to temperature and precipitation changes as well as the importance of temperature and precipitation for 213 species with estimates at ten or more grid cells, we adopted the within-subject centring approach^45^ under a hierarchical modelling framework to explicitly distinguish species-level effects (explaining variations in species-level responses between species) and population-level effects (explaining variations in population-level responses within species) of explanatory variables. Here we defined each species responses at their absolute latitudinal range mid-points as species-level responses, and responses within each grid cell as population-level responses.

In this model the species effect *μ*_*s*_, representing the species-level responses to temperature or precipitation increases in species *s*, is drawn from a normal distribution with mean of *ν*_*s*_ and variance of *σ* _*ν*_ ^*2*^. *ν*_*s*_ is further modelled with species-level explanatory variables:

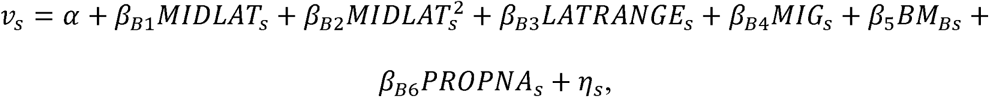

where *α* is the global intercept and *β*_*B1-B6*_ represent the species-level effects. *MIDLAT*_*s*_, *LATRANGE*_*s*_, *MIG*_*s*_, *BM*_*s*_, *PROPNA*_*s*_ are species-level explanatory variables; absolute latitudinal range mid-points, absolute latitudinal geographical range (degree), migration status (migrant or non-migrant), body mass (g, log_10_-transformed) and the mean proportion of missing values (i.e., interpolated values) in count records across all sites (%) for species *s*, respectively. *η*_*s*_ is a random term that accounts for phylogenetic dependence among species and is drawn from a multivariate normal distribution^46,47^:

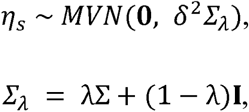

where Σ is a scaled variance-covariance matrix calculated from an ultrametric phylogenetic tree (defined below). By scaling Σ to a height of one, we can interpret *δ*^*2*^ as the residual variance^46^. For the strength of phylogenetic signal to vary, we also incorporated Pagel’s *λ*^48,49^ into the matrix with the identity matrix **I**. Here *λ* is a coefficient that multiplies the off-diagonal elements of Σ and a *λ* close to zero implies that the phylogenetic signal in the data is low, suggesting independence in the error structure of the data points, whereas a *λ* close to one suggests a good agreement with the Brownian Motion evolution model and thus suggests correlation in the error structure^46,49^. To incorporate uncertainties^50^ in phylogenetic trees in the calculation of Σ, we used a sample of 100 trees from a comprehensive avian phylogeny^51^ as the prior distribution for our analysis^46^. More specifically, one of the 100 trees was randomly drawn in each iteration and used for the calculation of Σ.

The population-level responses to temperature or precipitation increases *r*_*s,i*_ of species *s* in grid cell *i* was then assumed to derive from a normal distribution with mean *μ*_*s,i*_ and variance *σ* _*μ*_^*2*^, where *μ*_*s,i*_ is modelled using the species effect *μ*_*s*_:

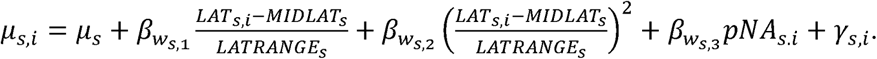

Here *β*_*Ws,1-3*_ represents the population-level effect of absolute latitudes *LAT*_*s,i*_ (in the form of linear and quadratic terms, to test non-linear patterns) and the mean proportion of missing values (i.e., interpolated values) in count records across all sites *pNA*_*s,i*_ (%) of grid cell *i* for species *s*. Here within-species variations in population-level responses (*μ*_*s,i*_ − *μ*_*s*_) are explained by within-species variations in absolute latitudes (*LAT*_*s,i*_ − *MIDLAT*_*s*_), divided by the absolute latitudinal geographical range of each species *LATRANGE*_*s*_, so that the estimated effects of absolute latitudes are comparable among species with varying latitudinal range size. The species-specific *β*_*Ws,1-3*_ is the random effect each governed by hyper-parameters as:

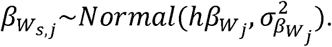

*γ*_*s,i*_ accounts for spatial autocorrelation within each species and is drawn from an intrinsic Gaussian conditional autoregressive (CAR) prior distribution with variance 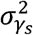:

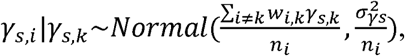

where *w*_*i,k*_ = 1 if grid cells *i* and *k* are neighbours, and 0 otherwise. *n*_*i*_ is the total number of neighbours of grid cell *i* and neighbours here are defined as those grid cells directly adjacent, including those diagonal. *σ*_*γs*_ ^*2*^ controls the amount of variation between the random effects.

We tested latitudinal patterns in the importance of temperature and precipitation using essentially the same model but the population-level importance of temperature or precipitation *imp*_*s,i*_ of species *s* in grid cell *i* was assumed to derive from a beta distribution with mean *c*_*s,i*_ and variance 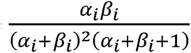 with a logit link function:

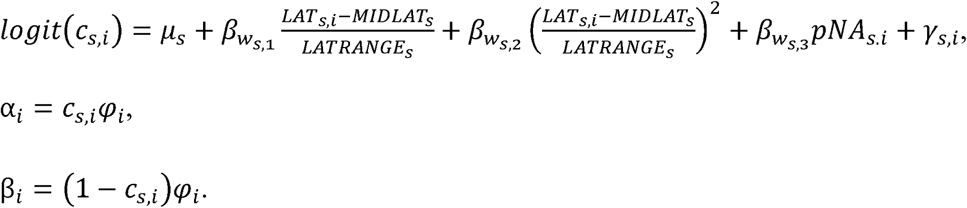

The models were implemented with OpenBUGS 3.2.3^52^ and the R2OpenBUGS package^53^ in R 3.4.1^41^. As non-informative prior distributions, we used a Gamma distribution with mean of 1 and variance of 100 for *φ*_*i*_ and the inverse of 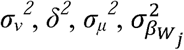 and 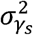, a uniform distribution on the interval [0, 1] for *λ*, normal distributions with mean of 0 and variance of 100 for *α, β*_*Bk*_, and *hβ*_*wj*_ Each MCMC algorithm was run with three chains with different initial values for 30,000 iterations with the first 10,000 discarded as burn-in and the remainder thinned to one in every four iterations to save storage space. Model convergence was checked with R-hat values.

Due to differences in the definition of species between the two sources used^51,54^, we combined two separate species defined in the BirdLife Checklist^54^ into one in four cases for this species-level analysis: Kentish plover *Charadrius alexandrinus* and snowy plover *C. nivosus*, common snipe *Gallinago gallinago* and Wilson’s snipe *G. delicata*, European herring gull *Larus argentatus* and Arctic herring gull *L. smithsonianus*, and common moorhen *Gallinula chloropus* and common gallinule *G. galeata. Larus glaucoides thayeri* was excluded from the latitudinal analysis as it is not included in either database. We also excluded from the analysis eight seabird species in Alcidae and Sulidae as neither the IWC nor CBC necessarily targets seabird species.

We also used R packages ape^55^, data.table^56^, dplyr^57^, ggplot2^58^, gridExtra^59^, mapdata^60^, plyr^61^, png^62^, RcolorBrewer^63^, rgdal^64^, raster^65^ and viridis^66^.

## Data Availability

The waterbird count data used in this study are collated and managed by Wetlands International and the National Audubon Society, and are available from Wetlands International at: http://iwc.wetlands.org/. The estimated abundance responses to temperature and precipitation as well as the importance of temperature and precipitation for each grid cell for each species are available as Supplementary Data S2. All the data on explanatory variables are freely available as specified in Supplementary Table S2.

## Code Availability

All the R codes used for the analyses are available as Supplementary Data S5-7.

## Acknowledgements

We thank the coordinators, thousands of volunteer counters and funders of the International Waterbird Census and Christmas Bird Count. T.A. was supported by the Grantham Foundation for the Protection of the Environment, the Kenneth Miller Trust and the Australian Research Council Future Fellowship (FT180100354). T.S. was funded by a Royal Society Wolfson Merit Award (WM170050) and by the National Research, Development and Innovation Office of Hungary (ÉLVONAL KKP-126949, K-116310). H.S.W. was supported by the Cambridge Trust Cambridge-Australia Poynton Scholarship and the Cambridge Department of Zoology JS Gardiner Fellowship. W.J.S. is supported by Arcadia and The David and Claudia Harding Foundation. This work is also funded by EU Horizon 2020 BACI project (Grant Agreement 640176), Ministry of the Environment of Japan, Environment Canada, AEWA Secretariat, EU LIFE+ NGO Operational Grant, MAVA Foundation, Swiss Federal Office for Environment and Nature, French Ministry of Environment and Sustainable Development, UK Department of Food and Rural Affairs, Norwegian Nature Directorate, Dutch Ministry of Economics, Agriculture and Innovation, DOB Ecology and Wetlands International members. Thanks to M. Amano for all the support.

## Author contributions

T.A. designed the study. T.A., T.S., H.S.W., B.S., S.N., T.M., T.L., D.B. and N.L.M. collected and prepared data for the analyses. T.A. analysed the data and wrote the paper. All authors discussed the results and commented on the manuscript at all stages.

## Competing interests

The authors declare no competing interests.

## Additional information

### Supplementary information

is available for this paper.

