## Supplementary Information for "Responses of global waterbird populations to climate change vary with latitude"

**Supplementary Table S1. Hypotheses tested for explaining among- and within-species latitudinal variations in waterbird abundance responses to temperature and precipitation changes.**

| Hypotheses | | Expected latitudinal patterns (each dot represents the range mid-point of each species, with solid lines showing within-species patterns; dotted lines represent zero, i.e., no response). Note that latitudes are absolute values; see Methods for more detail. | Supporting evidence |
|---|---|---|---|
| Abundance changes with increasing temperature | Among species | Monotonic increase with absolute latitudes, with negative responses in species at lower latitudes 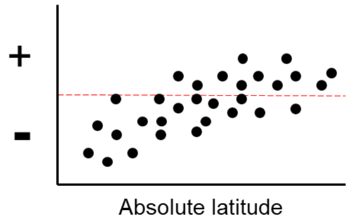  | Vulnerability to warming temperature increases from polar towards tropical regions as species in the tropics tend to live closer to their upper temperature limits[15] |
| | Within species | Monotonic increase with absolute latitudes, with negative responses in populations at lower latitudes and positive responses in populations at higher latitudes 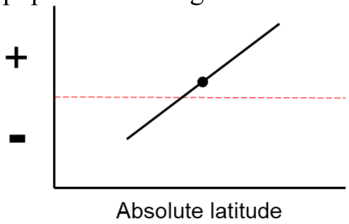  | Populations at higher latitudes benefit from, and those at lower latitudes are negatively affected by, temperature increase[67], if all populations have the same fitness curve with the optimal climatic niche at the latitudinal midpoint of its range[68] |
| Importance of temperature in explaining abundance changes | Among species | Monotonic increase with absolute latitudes 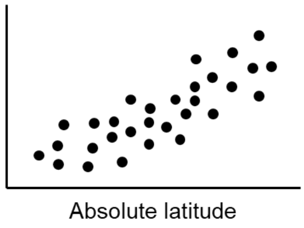  | Species-richness patterns at high latitudes are often limited by ambient energy (correlated with temperature) and at low latitudes by moisture availability[28]. If species richness represents the summation of individual population response, the same is applicable to species-level responses in abundance[4]. |
| | Within species | Higher at both range edges | If climatic factors limit species distributions[69], they are expected to be more important determinants of species abundance at range edges[12]. |

| | | | |
|---|---|---|---|
| | | 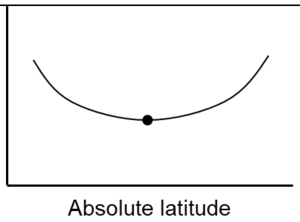  | |
| Abundance changes with increasing precipitation | Among species | Monotonic decrease, with negative responses in species at higher latitudes 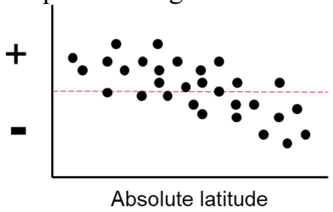  | Increase in Dec-Feb precipitation (and associated increase in water level, decrease in habitat heterogeneity, and increased thermoregulatory cost under wet weather) is observed to negatively affect waterbird abundance at higher latitudes (e.g., Europe[70,71] and Argentina[32]) while higher precipitation increases the availability of wetlands and vegetation in dry parts of the tropics (e.g., the Sahel region[31,22]) although effects may depend on each species' ecology[22]. |
| | Within species | Monotonic decrease, with negative responses in populations at higher latitudes 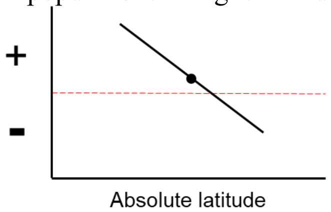  | Within-species patterns are expected to be similar to among-species patterns, especially for wide-ranging species. |
| Importance of precipitation in explaining abundance changes | Among species | Monotonic decrease with absolute latitudes 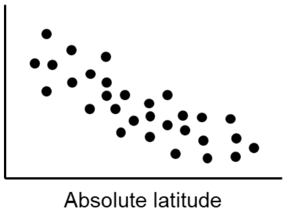  | Species-richness patterns at high latitudes are often limited by ambient energy and at low latitudes by moisture availability (correlated with precipitation)[28]. Assuming that species richness represents the summation of individual population response, the same is applicable to species-level responses in abundance[4]. |
| | Within species | Higher at both range edges 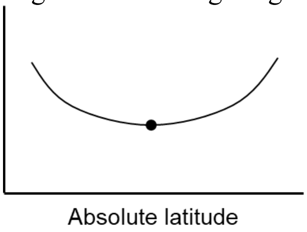  | If climatic factors limit species distributions[69], they are expected to be more important determinants of species abundance at range edges[12]. |

**Supplementary Table S2. Additional hypotheses tested for explaining among-species variations in waterbird abundance responses to temperature and precipitation changes.**

| Hypotheses | Expected patterns | Variables used | Data sources |
|---|---|---|---|
| Latitudinal geographical range | • Species with a narrower latitudinal range have narrower temperature niche, thus more vulnerable to temperature increases (i.e., the rate of abundance changes with increasing temperature is more negative)[72].<br>• Species with a narrower latitudinal range have narrower temperature niche, thus their abundance is affected more by temperature changes (i.e., the importance of temperature is higher)[72]. | Differences between maximum and minimum absolute latitudes of geographical range | BirdLife Data Zone* |
| Migratory status | • Resident species can be more negatively affected by temperature increases, due to their limited dispersal ability (i.e., the rate of abundance changes with increasing temperature is more negative)[25].<br>• Migratory species generally have a higher dispersal ability[25] and track climate niches to a greater extent than resident species[26], thus can be more responsive to changes in local temperature and precipitation (i.e., the importance of temperature and precipitation is higher).<br>• Migratory species often show fidelity to breeding and non-breeding sites between years, thus may be less responsive to changes in local temperature and precipitation (i.e., the importance of temperature and precipitation is lower)[73].<br>• Migratory species can also be affected by conditions at multiple locations, thus local climatic conditions may play a limited role in explaining their abundance (i.e., the importance of temperature and precipitation is lower)[74]. | Migrant or non-migrant | BirdLife Data Zone* |
| Body size | • Smaller-sized species can be more negatively affected by increasing temperature, due to their limited dispersal ability (i.e., the rate of abundance changes with increasing temperature is more negative)[25].<br>• Larger-sized species have a higher dispersal ability, thus may be more responsive to changes in local temperature and precipitation | Body mass (g) | Elton Traits 1.0[75] |

| | |
|---|---|
| | (i.e., the importance of temperature and precipitation is higher)[25]. |

* http://datazone.birdlife.org/home

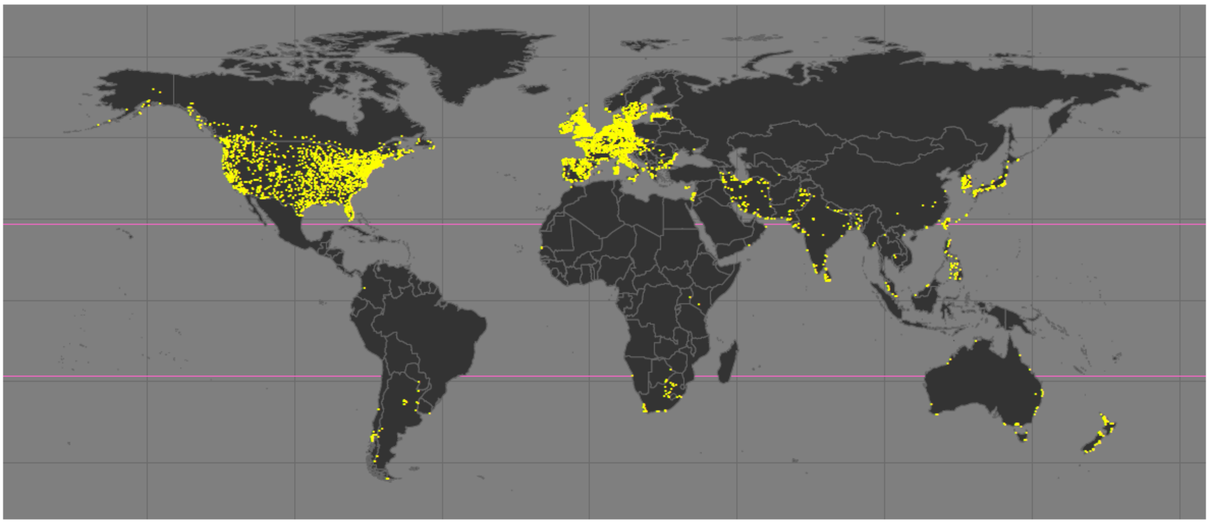

**Supplementary Figure S1. Distribution of the 6,822 survey sites used in the analyses.** The area between pale pink lines represents the tropical region.

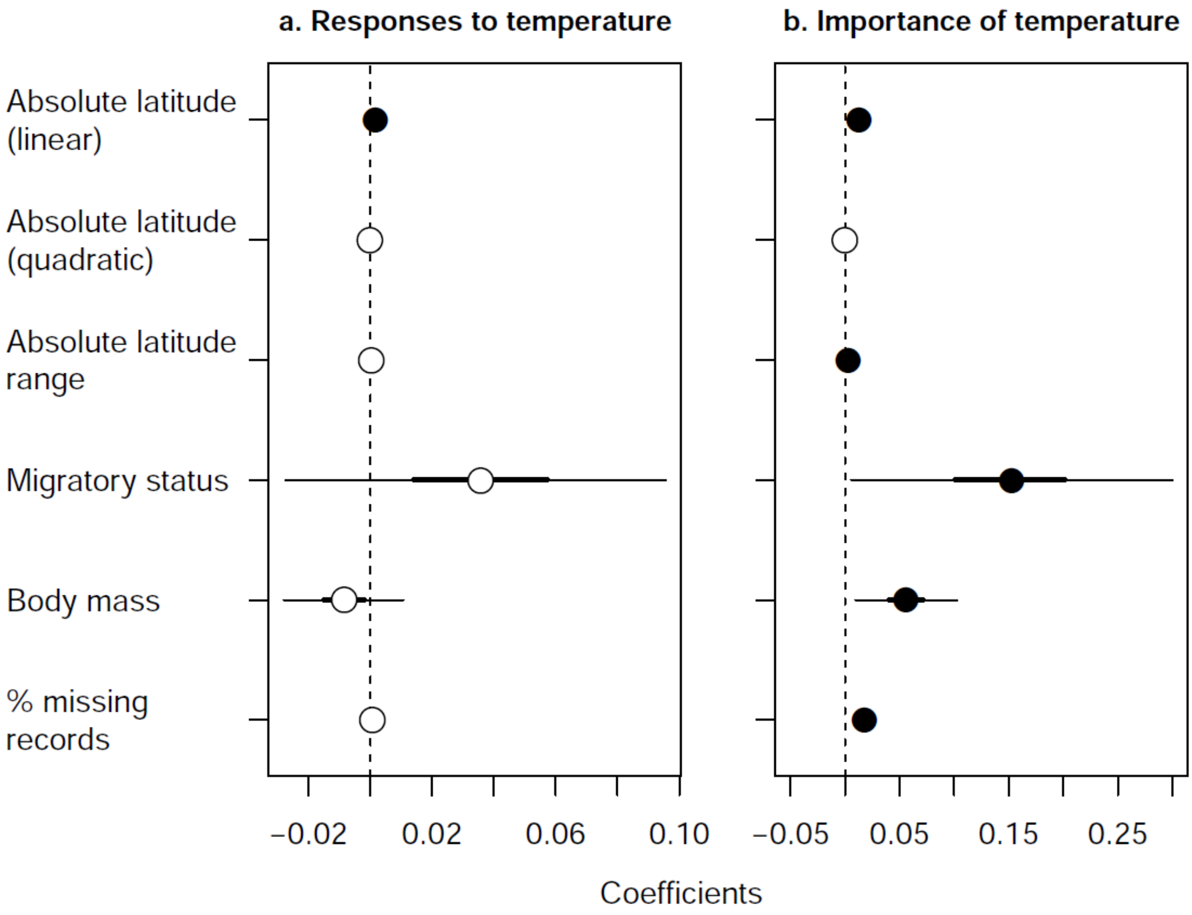

**Supplementary Figure S2. Effects of species-level predictors on waterbird abundance responses to temperature changes.** The estimated coefficients with 95% and 50% (thick lines) credible intervals of six explanatory variables for explaining among-species variations in the rate of abundance changes with increasing temperature (a) and the importance of temperature in explaining abundance changes (b). Filled circles indicate variables with 95% credible intervals not overlapping with zero.

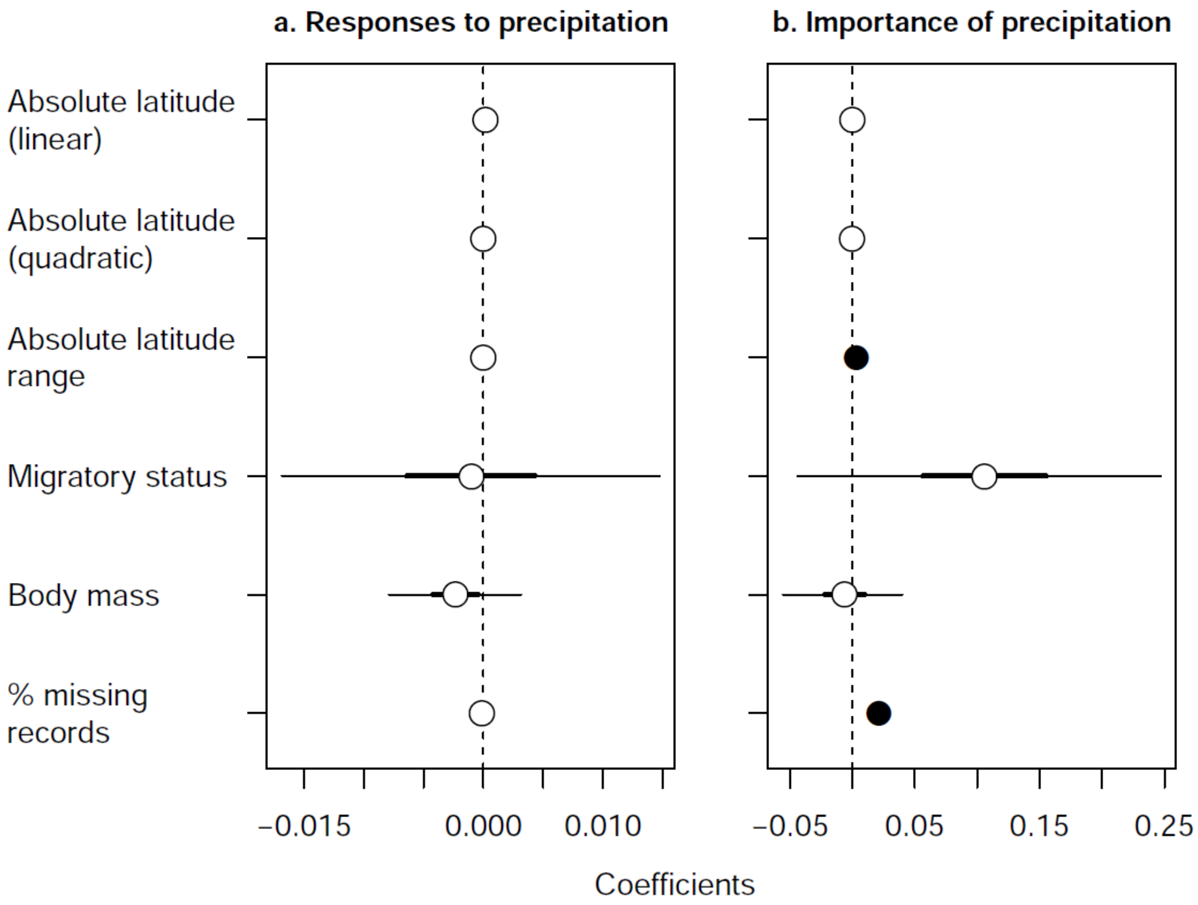

**Supplementary Figure S3. Effects of species-level predictors on waterbird abundance responses to precipitation changes.** The estimated coefficients with 95% and 50% (thick lines) credible intervals of six explanatory variables for explaining among-species variations in the rate of abundance changes with increasing precipitation (a) and the importance of precipitation in explaining abundance changes (b). Filled circles indicate variables with 95% credible intervals not overlapping with zero.

**Supplementary Data**

**Supplementary Data S1** (available from the corresponding author upon request)
Species-level maps of distribution of estimated abundance responses to changes in temperature and precipitation in each 1° × 1° grid cell. The rate of abundance changes with increasing temperature (a), the independent capacity of temperature in explaining abundance changes (b), the rate of abundance changes with increasing precipitation (c) and the independent capacity of precipitation in explaining abundance changes (d). The region between the yellow solid lines is the tropics. Species names are based on the BirdLife Checklist[54].

**Supplementary Data S2** (separate file)
Estimated abundance responses to changes in temperature and precipitation in each 1° × 1° grid cell (*Grid* with its coordinates shown in *Lon_Grid* and *Lat_Grid*) for each of the 390 species (*BLsciname* based on the BirdLife Checklist[54]). The rate of abundance changes with increasing temperature (*mnTcoef*) with its standard error (*seTcoef*), the rate of abundance changes with increasing precipitation (*mnPcoef*) with its standard error (*sePcoef*), the rate of abundance changes with abundance in the previous year (*mnARcoef*) with its standard error (*seARcoef*), the independent capacity of temperature in explaining abundance changes (*mnTind*), the independent capacity of precipitation in explaining abundance changes (*mnPind*), the independent capacity of abundance in the previous year in explaining abundance changes (*mnARind*) and the mean proportion of missing values (i.e., interpolated values) in count records across all sites (*mnNAprop*).

**Supplementary Data S3** (separate file)
The posterior median coefficients (columns coded with *50%*) with 95% credible intervals (*2.5%*, *97.5%*) of three population-level predictors (*Lat-linear*, *Lat-quadratic*, *NA*) for explaining within-species variations in the rate of waterbird abundance changes with increasing temperature (*Tcoef*), the importance of temperature in explaining abundance changes (*Tcont*), the rate of abundance changes with increasing precipitation (*Pcoef*) and the importance of precipitation in explaining abundance changes (*Pcont*). Values are shown only for the 213 species for which there were estimates at ten or more grid cells and thus were analysed for latitudinal pattenrs (*BLsciname* based on the BirdLife Checklist[54]).

**Supplementary Data S4** (separate file)

A list of the 390 waterbird species analysed in this study. The explanations and references of column names are as follows: *BLsciname*: scientific name, *BLengname*: English name, *BLfamily*: family name, *BLgroup*: taxonomic groups defined by the BirdLife International[54], *IOCGroups*: taxonomic groups defined by the International Ornithological Congress[39], *PHYsciname*: scientific name used in phylogenetic trees, *PHYlabel*: labels used in phylogenetic trees[51], *SISRecID*: unique IDs assigned to each taxonomic entity[54], *BodyMass*: body mass (g)[75], *mig.status*: migration status, *maxlat*: the maximum latitude of geographical range, *minlat*: the minimum latitude of geographical range, *midlat*: the latitudinal mid-point of geographical range, *absmaxlat*: the maximum absolute latitude of geographical range, *absmidlat*: the absolute latitudinal mid-point of geographical range, *absminlat*: the minimum absolute latitude of geographical range (http://datazone.birdlife.org/home), *muP*: parameter p to determine the relationship between survey effort and the number of birds counted, *muB*: parameter B to determine the relationship between survey effort and the number of birds counted[16].

**Supplementary Data S5** (separate file)

The R script for estimating abundance responses to temperature and precipitation changes and the importance of temperature and precipitation.

**Supplementary Data S6** (separate file)

The R script for analysing among- and within-species latitudinal variations in abundance responses to temperature and precipitation changes.

**Supplementary Data S7** (separate file)

The R script for analysing among- and within-species latitudinal variations in the importance of temperature and precipitation.
