## Supplementary Data S5-7 for "Responses of global waterbird populations to climate change vary with latitude": SupplementaryDataS5_Model_AbundanceResponses.r

```
## R script for estimating abundance responses to temperature and precipitation changes
## all data files are available from the corresponding author except for 
## Modelleddata.all.csv, which is available upon request at http://iwc.wetlands.org/

rm(list=ls())

## upload packages ---------------------------
library(plyr)
library(dplyr)
library(data.table)
library(hier.part)
library(zoo)

setwd("")


## climate data ----------------------------------
## SitesClimate_tmp.csv and SitesClimate_pre.csv are data on mean temperature and precipitation for each site in each year
tmp <- fread("SitesClimate_tmp.csv")
tmp$Year <- as.numeric(tmp$Year)
pre <- fread("SitesClimate_pre.csv")
pre$Year <- as.numeric(pre$Year)
colnames(tmp)[5:16] <- colnames(pre)[5:16] <- c("Jan","Feb","Mar","Apr","May","Jun","Jul","Aug","Sep","Oct","Nov","Dec")


## Species data -------------------------------------
## Modelleddata.all.csv is raw data (available upon request at http://iwc.wetlands.org/)
Alldata <- fread("Modelleddata.all.csv")

# site data: SiteSpeciesDataFrame.csv is information on each survey site with observed species
spst <- fread("SiteSpeciesDataFram.csv")
spst <- spst[spst$NumYearsSurveyed>9, ] # only sites with 10 or more non-zero records since 1990

# use only species with at least one site with 10 or more non-zero records since 1990
# note species name is largely IOCsciname with 4 exceptions
SPname <- sort(unique(spst$Species))
Nsp <- length(SPname)
# known issues with species name
# Anas poecilorhyncha (i=28) -> Anas poecilorhyncha and Anas zonorhyncha (Spot-billed duck)
# Charadrius aquilonius (i=104) -> Charadrius obscurus (New Zealand Plover)
# Larus glaucoides (i=236) -> Larus glaucoides_thayeri (Iceland_Thayer's Gull)
# Rallus crepitans (i=336) -> Rallus crepitans and Rallus obsoletus (Clapper Rail and Ridgway's Rail)
SPIOCname <- SPname
SPIOCname[28] <- "Anas poecilorhyncha and Anas zonorhyncha"
SPIOCname[104] <- "Charadrius obscurus"
SPIOCname[236] <- "Larus glaucoides_thayeri"
SPIOCname[336] <- "Rallus crepitans and Rallus obsoletus"

# For CBC species, read effort parameters
CBCsp <- read.csv("CBCsp_PB.csv")


### define a function applied to each site (tmp and pre as predictors) ------------------
TPcontribution2 <- function(x){
  tdata <- x
  
  # read temp and pre data for the site
  stmp <- tmp[tmp$Site==tdata$Site[1],]
  spre <- pre[pre$Site==tdata$Site[1],]
  
  # for CBC site read effort parameters
  if(tdata[nrow(tdata),"Scheme"]=="CBC"){
    muP <- as.numeric(CBCsp[CBCsp$IOCsciname==tdata[nrow(tdata),"IOCsciname"],"muP"])
    muB <- as.numeric(CBCsp[CBCsp$IOCsciname==tdata[nrow(tdata),"IOCsciname"],"muB"])
  }
  
  # restrict only to post-1990 data
  years <- data.frame(Year=max(1990,min(tdata$Year)):max(tdata$Year))
  tdata <- merge(years, tdata, by="Year", all.x=T)
  tdata <- tdata[which(!is.na(tdata$Count))[1]:nrow(tdata),] # starting from the first year with a record (e.g., if no record in 1990, remove 1990) 
  tdata$Count <- na.approx(tdata$Count)  # missing value imputation based by linear interpolation
  tdata$YearPrev1 <- tdata$Year-1
  tdata$YearPrev2 <- tdata$Year-2
  
  # only if there are 10 or more records AND 5 or more non-zero records (excluding imputed records) after 1990
  (
  if((length(which(tdata[!is.na(tdata$Site),"Count"]>0))>4)&&(nrow(tdata[!is.na(tdata$Site),])>=10)){
  
    # climate data in Dec Year t-2 for IWC data
    tdata <- merge(tdata, stmp[,c("Year","Dec")], by.x="YearPrev2", by.y="Year")
    colnames(tdata)[ncol(tdata)] <- "TmpDecPrev2"
    tdata <- merge(tdata, spre[,c("Year","Dec")], by.x="YearPrev2", by.y="Year")
    colnames(tdata)[ncol(tdata)] <- "PreDecPrev2"
    
    # climate data in Jan, Feb and Dec Year t-1
    tdata <- merge(tdata, stmp[,c("Year","Jan","Feb","Dec")], by.x="YearPrev1", by.y="Year")
    colnames(tdata)[(ncol(tdata)-2):ncol(tdata)] <- c("TmpJanPrev1","TmpFebPrev1","TmpDecPrev1")
    tdata <- merge(tdata, spre[,c("Year","Jan","Feb","Dec")], by.x="YearPrev1", by.y="Year")
    colnames(tdata)[(ncol(tdata)-2):ncol(tdata)] <- c("PreJanPrev1","PreFebPrev1","PreDecPrev1")
    
    # climate data in Jan, Feb and Dec Year t
    tdata <- merge(tdata, stmp[,c("Year","Jan","Feb","Dec")], by="Year")
    colnames(tdata)[(ncol(tdata)-2):ncol(tdata)] <- c("TmpJan","TmpFeb","TmpDec")
    tdata <- merge(tdata, spre[,c("Year","Jan","Feb","Dec")], by="Year")
    colnames(tdata)[(ncol(tdata)-2):ncol(tdata)] <- c("PreJan","PreFeb","PreDec")
    
    # climate at previous wintering season
    (if(tdata[nrow(tdata),"Scheme"]!="CBC"){
      tdata$TmpPrev <- apply(tdata[,c("TmpDecPrev2","TmpJanPrev1","TmpFebPrev1")], 1, mean)
      tdata$PrePrev <- apply(tdata[,c("PreDecPrev2","PreJanPrev1","PreFebPrev1")], 1, mean)
      tdata$TmpSame <- tdata$TmpDecPrev1
      tdata$PreSame <- tdata$PreDecPrev1
    }
    else{ # if CBC
      tdata$TmpPrev <- apply(tdata[,c("TmpDecPrev1","TmpJan","TmpFeb")], 1, mean)
      tdata$PrePrev <- apply(tdata[,c("PreDecPrev1","PreJan","PreFeb")], 1, mean)
      tdata$TmpSame <- tdata$TmpDec
      tdata$PreSame <- tdata$PreDec
    })
    
    # count at Year t-1
    tdata$CountPrev <- c(NA, tdata$Count[1:(nrow(tdata)-1)]) 
    # remove the first row (NA for count at Year t-1)
    tdata <- tdata[-1,]
    
    # only if there are cimate data for the site
    (
    if(nrow(na.omit(tdata[,c("TmpPrev","PrePrev","TmpSame","PreSame")]))>9){
      ## modelling
      (
        if(tdata[nrow(tdata),"Scheme"]!="CBC"){ # non CBC
          # temp model
          TmodelPrev <- lm((log(Count+0.5) - log(CountPrev+0.5)) ~ TmpPrev + log(CountPrev+0.5), data=tdata) # with climate at the previous season
          TmodelSame <- lm((log(Count+0.5) - log(CountPrev+0.5)) ~ TmpSame + log(CountPrev+0.5), data=tdata) # with climate at the same season
          TID <- 1  # 1=Tmp at the previous season
          if(AIC(TmodelPrev) > AIC(TmodelSame)) TID <- 2 # 2=Tmp at the same season
          
          # prec model
          PmodelPrev <- lm((log(Count+0.5) - log(CountPrev+0.5)) ~ PrePrev + log(CountPrev+0.5), data=tdata) # with climate at the previous season
          PmodelSame <- lm((log(Count+0.5) - log(CountPrev+0.5)) ~ PreSame + log(CountPrev+0.5), data=tdata) # with climate at the same season
          PID <- 1 # 1=Pre at the previous season
          if(AIC(PmodelPrev) > AIC(PmodelSame)) PID <- 2 # 2=Pre at the same season
          
          pred <- data.frame(TmpPrev=tdata$TmpPrev, TmpSame=tdata$TmpSame, PrePrev=tdata$PrePrev, PreSame=tdata$PreSame)
          hp_res <- hier.part((log(tdata$Count+0.5) - log(tdata$CountPrev+0.5)), 
                              data.frame(Tmp=pred[,TID], Pre=pred[,2+PID], logPrevCount=log(tdata$CountPrev+0.5)),
                              family="gaussian", gof="Rsqu", barplot=F)
          
          fmodel <- lm((log(tdata$Count+0.5) - log(tdata$CountPrev+0.5)) ~ pred[,TID] + pred[,2+PID] + log(tdata$CountPrev+0.5))
        }
        else{ # CBC
          # temp model
          TmodelPrev <- lm((log(Count+0.5) - log(CountPrev+0.5)) ~ TmpPrev + log(CountPrev+0.5), data=tdata,
                           offset=muB*((tdata$Hours/mean(tdata$Hours, na.rm=T))^muP-1)/muP) # with climate at the previous season
          TmodelSame <- lm((log(Count+0.5) - log(CountPrev+0.5)) ~ TmpSame + log(CountPrev+0.5), data=tdata,
                           offset=muB*((tdata$Hours/mean(tdata$Hours, na.rm=T))^muP-1)/muP) # with climate at the same season
          TID <- 1  # 1=Tmp at the previous season
          if(AIC(TmodelPrev) > AIC(TmodelSame)) TID <- 2 # 2=Tmp at the same season
          
          # prec model
          PmodelPrev <- lm((log(Count+0.5) - log(CountPrev+0.5)) ~ PrePrev + log(CountPrev+0.5), data=tdata,
                           offset=muB*((tdata$Hours/mean(tdata$Hours, na.rm=T))^muP-1)/muP) # with climate at the previous season
          PmodelSame <- lm((log(Count+0.5) - log(CountPrev+0.5)) ~ PreSame + log(CountPrev+0.5), data=tdata,
                           offset=muB*((tdata$Hours/mean(tdata$Hours, na.rm=T))^muP-1)/muP) # with climate at the same season
          PID <- 1 # 1=Pre at the previous season
          if(AIC(PmodelPrev) > AIC(PmodelSame)) PID <- 2 # 2=Pre at the same season
          
          pred <- data.frame(TmpPrev=tdata$TmpPrev, TmpSame=tdata$TmpSame, PrePrev=tdata$PrePrev, PreSame=tdata$PreSame)
          hp_res <- hier.part((log(tdata$Count+0.5)-log(tdata$CountPrev+0.5)-muB*((tdata$Hours/mean(tdata$Hours, na.rm=T))^muP-1)/muP), 
                              data.frame(Tmp=pred[,TID], Pre=pred[,2+PID], logPrevCount=log(tdata$CountPrev+0.5)),
                              family="gaussian", gof="Rsqu", barplot=F)
          
          fmodel <- lm((log(tdata$Count+0.5) - log(tdata$CountPrev+0.5)) ~ pred[,TID] + pred[,2+PID] + log(tdata$CountPrev+0.5),
                       offset=muB*((tdata$Hours/mean(tdata$Hours, na.rm=T))^muP-1)/muP)
        }
      )
      data.frame(IOCsciname=tdata[nrow(tdata),"IOCsciname"], IOCengname=tdata[nrow(tdata),"IOCengname"],
                 BLsciname=tdata[nrow(tdata),"BLsciname"], BLengname=tdata[nrow(tdata),"BLengname"],
                 Lon_Site=tdata[nrow(tdata),"Lon_Site"], Lat_Site=tdata[nrow(tdata),"Lat_Site"], Grid=tdata[nrow(tdata),"Grid"],
                 Lon_Grid=tdata[nrow(tdata),"Lon_Grid"], Lat_Grid=tdata[nrow(tdata),"Lat_Grid"], Scheme=tdata[nrow(tdata),"Scheme"],
                 N=nrow(tdata), N_nonzero=nrow(tdata[tdata$Count>0,]), N_NA=length(which(is.na(tdata$Site))),
                 Tvar=as.character(colnames(pred)[TID]), Pvar=as.character(colnames(pred)[2+PID]),
                 Tcoef=round(summary(fmodel)$coefficients[2, "Estimate"], digits=5),
                 Pcoef=round(summary(fmodel)$coefficients[3, "Estimate"], digits=5),
                 ARcoef=round(summary(fmodel)$coefficients[4, "Estimate"], digits=5),
                 Tse=round(summary(fmodel)$coefficients[2, "Std. Error"], digits=5),
                 Pse=round(summary(fmodel)$coefficients[3, "Std. Error"], digits=5),
                 ARse=round(summary(fmodel)$coefficients[4, "Std. Error"], digits=5),
                 Tind=round(hp_res$IJ[,"I"], digits=5)[1], Pind=round(hp_res$IJ[,"I"], digits=5)[2], ARind=round(hp_res$IJ[,"I"], digits=5)[3]
                 )
    }
    else{
      data.frame(IOCsciname=tdata[nrow(tdata),"IOCsciname"], IOCengname=tdata[nrow(tdata),"IOCengname"],
                 BLsciname=tdata[nrow(tdata),"BLsciname"], BLengname=tdata[nrow(tdata),"BLengname"],
                 Lon_Site=tdata[nrow(tdata),"Lon_Site"], Lat_Site=tdata[nrow(tdata),"Lat_Site"], Grid=tdata[nrow(tdata),"Grid"],
                 Lon_Grid=tdata[nrow(tdata),"Lon_Grid"], Lat_Grid=tdata[nrow(tdata),"Lat_Grid"], Scheme=tdata[nrow(tdata),"Scheme"],
                 N=nrow(tdata), N_nonzero=nrow(tdata[tdata$Count>0,]), N_NA=length(which(is.na(tdata$Site))),
                 Tvar="NoData", Pvar="NoData",
                 Tcoef=NA, Pcoef=NA, ARcoef=NA, Tse=NA, Pse=NA, ARse=NA, Tind=NA, Pind=NA, ARind=NA
      )
    }
    )
  }
  else{ # if there are not enough records at the site
    data.frame(IOCsciname=tdata[nrow(tdata),"IOCsciname"], IOCengname=tdata[nrow(tdata),"IOCengname"],
               BLsciname=tdata[nrow(tdata),"BLsciname"], BLengname=tdata[nrow(tdata),"BLengname"],
               Lon_Site=tdata[nrow(tdata),"Lon_Site"], Lat_Site=tdata[nrow(tdata),"Lat_Site"], Grid=tdata[nrow(tdata),"Grid"],
               Lon_Grid=tdata[nrow(tdata),"Lon_Grid"], Lat_Grid=tdata[nrow(tdata),"Lat_Grid"], Scheme=tdata[nrow(tdata),"Scheme"],
               N=nrow(tdata), N_nonzero=nrow(tdata[tdata$Count>0,]), N_NA=length(which(is.na(tdata$Site))),
               Tvar="NA", Pvar="NA",
               Tcoef=NA, Pcoef=NA, ARcoef=NA, Tse=NA, Pse=NA, ARse=NA, Tind=NA, Pind=NA, ARind=NA
    )
  }
  )
}


### define a function applied to each species (tmp and pre as predictors) ----------------------
TPcont.to.sp <- function(x){
  sdata <- x
  SPID <- which(SPIOCname==as.character(sdata[1,"IOCsciname"]))
  sst <- spst[spst$Species==SPname[SPID],]
  
  # only sites with 10 or more non-zero records since 1990
  Efdata <- filter(sdata, Site %in% sst$Site)
  
  # Apply analysis to each site
  ddply(Efdata, .(Site), TPcontribution2)
}


### apply defined analysis to each species's data with ddply (tmp and pre as predictors) -------------------------------
# extract only data with species for analysis
sAlldata <- filter(Alldata, IOCsciname %in% SPIOCname)

start <- Sys.time()
datalist <-  list()
TPres2 <- ddply(sAlldata, .(IOCsciname), TPcont.to.sp)
print(Sys.time()-start)

write.csv(TPres2, "TPcontribution.all.csv", row.names=F)


### collating results at the grid level (tmp and pre) ----------------

res <- fread("TPcontribution.all.csv")
# remove NAs
res <- na.omit(res)
# remove results with 9 or fewer non-zero observations
res <- res[res$N_nonzero>9,]
# IOCsciname="Larus glaucoides_thayeri" does not have BLsciname, so use IOCsiname
res[res$BLsciname=="", "BLsciname"] <- "Larus glaucoides_thayeri"


## define a function to calculate grid-level stats
SitetoGrid <- function(d){
  data.frame(Lon_Grid=d$Lon_Grid[1], Lat_Grid=d$Lat_Grid[1], N=nrow(d),
             mnTcoef=sum(d$Tcoef/(d$Tse)^2)/sum(1/(d$Tse)^2),
             seTcoef=sqrt(1/sum(1/(d$Tse)^2)),
             mnPcoef=sum(d$Pcoef/(d$Pse)^2)/sum(1/(d$Pse)^2),
             sePcoef=sqrt(1/sum(1/(d$Pse)^2)),
             mnARcoef=sum(d$ARcoef/(d$ARse)^2)/sum(1/(d$ARse)^2),
             seARcoef=sqrt(1/sum(1/(d$ARse)^2)),
             mnTind=mean(d$Tind, na.rm = T),
             mnPind=mean(d$Pind, na.rm = T),
             mnARind=mean(d$ARind, na.rm = T),
             wmnTind=sum(d$Tind*d$N)/sum(d$N),
             wmnPind=sum(d$Pind*d$N)/sum(d$N),
             wmnARind=sum(d$ARind*d$N)/sum(d$N),
             mnNAprop=mean(d$N_NA/d$N, na.rm=T)
             )
  }
Gridsummary <- ddply(res, .(BLsciname, Grid), SitetoGrid) # based on BLsciname

## calculate community-level grid means
CalcComMean <- function(d){
  data.frame(Lon_Grid=d$Lon_Grid[1], Lat_Grid=d$Lat_Grid[1], Nsp=nrow(d),
            commnTcoef=sum(d$mnTcoef/(d$seTcoef)^2)/sum(1/(d$seTcoef)^2),
            comseTcoef=sqrt(1/sum(1/(d$seTcoef)^2)),
            commnPcoef=sum(d$mnPcoef/(d$sePcoef)^2)/sum(1/(d$sePcoef)^2),
            comsePcoef=sqrt(1/sum(1/(d$sePcoef)^2)),
            commnARcoef=sum(d$mnARcoef/(d$seARcoef)^2)/sum(1/(d$seARcoef)^2),
            comseARcoef=sqrt(1/sum(1/(d$seARcoef)^2)),
            commnTind=mean(d$mnTind, na.rm = T),
            commnPind=mean(d$mnPind, na.rm = T),
            commnARind=mean(d$mnARind, na.rm = T),
            comwmnTind=sum(d$wmnTind*d$N)/sum(d$N),
            comwmnPind=sum(d$wmnPind*d$N)/sum(d$N),
            comwmnARind=sum(d$wmnARind*d$N)/sum(d$N),
            commnNAprop=mean(d$mnNAprop, na.rm=T)
            )
}
Gridcommean <- ddply(Gridsummary, .(Grid), CalcComMean)

write.csv(Gridsummary, "Grid.population-level.mean.csv", row.names=F)
write.csv(Gridcommean, "Grid.community-level.mean.csv", row.names=F)
```
