## Supplementary Data S5-7 for "Responses of global waterbird populations to climate change vary with latitude": SupplementaryDataS6_LatitudinalAnalysis_Response.r

```
## R script for latitudinal analyses (abundance responses to temperature and precipitation)
## all data files are available from the corresponding author

rm(list=ls())

### upload packages ---------------------------
library(R2OpenBUGS)
library(ape)
library(plyr)

setwd("")


### prepare necessary data -----------------

## grid cell data
respvar <- read.csv("Grid.population-level.mean.csv")

# exclude species with 9 or fewer grids
SPincG <- names(which(table(respvar$BLsciname)>9)) # based on BLsciname
respvar <- respvar[respvar$BLsciname %in% SPincG, ]

# following species are not separated in BirdTree, so combined
# Charadrius alexandrinus AND nivosus
# Gallinago gallinago AND delicata
# Gallinula chloropus AND galeata
# Larus argentatus AND smithsonianus
respvar$BLsciname <- gsub('Charadrius nivosus', 'Charadrius alexandrinus', respvar$BLsciname)
respvar$BLsciname <- gsub('Gallinago delicata', 'Gallinago gallinago', respvar$BLsciname)
respvar$BLsciname <- gsub('Gallinula galeata', 'Gallinula chloropus', respvar$BLsciname)
respvar$BLsciname <- gsub('Larus smithsonianus', 'Larus argentatus', respvar$BLsciname)

# exclude "Larus glaucoides_thayeri" (not in BL and BirdTree)
respvar <- respvar[respvar$BLsciname!="Larus glaucoides_thayeri",]

## species data
SPinc <- names(table(respvar$BLsciname))
# information on latitudinal distribution ranges
rangeinfo <- read.csv("RangeMidPoints.csv")
# combine range information for the eight species with issues above
range1 <- rangeinfo[rangeinfo$BLsciname %in% c('Charadrius nivosus', 'Charadrius alexandrinus'),]
rangeinfo[rangeinfo$BLsciname=='Charadrius alexandrinus', 
          c("maxlat","minlat","midlat","absmaxlat","absmidlat","absminlat")] <- c(max(range1$maxlat), min(range1$minlat),
                                                                                  median(c(max(range1$maxlat), min(range1$minlat))),
                                                                                  max(range1$absmaxlat),
                                                                                  median(c(max(range1$absmaxlat),min(range1$absminlat))),
                                                                                  min(range1$absminlat))
range1 <- rangeinfo[rangeinfo$BLsciname %in% c('Gallinago delicata', 'Gallinago gallinago'),]
rangeinfo[rangeinfo$BLsciname=='Charadrius gallinago', 
          c("maxlat","minlat","midlat","absmaxlat","absmidlat","absminlat")] <- c(max(range1$maxlat), min(range1$minlat),
                                                                                  median(c(max(range1$maxlat), min(range1$minlat))),
                                                                                  max(range1$absmaxlat),
                                                                                  median(c(max(range1$absmaxlat),min(range1$absminlat))),
                                                                                  min(range1$absminlat))
range1 <- rangeinfo[rangeinfo$BLsciname %in% c('Gallinula galeata', 'Gallinula chloropus'),]
rangeinfo[rangeinfo$BLsciname=='Gallinula chloropus', 
          c("maxlat","minlat","midlat","absmaxlat","absmidlat","absminlat")] <- c(max(range1$maxlat), min(range1$minlat),
                                                                                  median(c(max(range1$maxlat), min(range1$minlat))),
                                                                                  max(range1$absmaxlat),
                                                                                  median(c(max(range1$absmaxlat),min(range1$absminlat))),
                                                                                  min(range1$absminlat))
range1 <- rangeinfo[rangeinfo$BLsciname %in% c('Larus smithsonianus', 'Larus argentatus'),]
rangeinfo[rangeinfo$BLsciname=='Larus argentatus', 
          c("maxlat","minlat","midlat","absmaxlat","absmidlat","absminlat")] <- c(max(range1$maxlat), min(range1$minlat),
                                                                                  median(c(max(range1$maxlat), min(range1$minlat))),
                                                                                  max(range1$absmaxlat),
                                                                                  median(c(max(range1$absmaxlat),min(range1$absminlat))),
                                                                                  min(range1$absminlat))

rangeinfo <- rangeinfo[rangeinfo$BLsciname %in% SPinc, ]
# species-level information
sp <- read.csv("Species-level.summary090117.traits.phylo.csv")
sp <- merge(rangeinfo[,c(1,3:8)], sp, by="BLsciname")

# exclude seabird species in Alcidae (8) and Sulidae (1 species)
sp <- sp[sp$BLfamily!="Alcidae",]
sp <- sp[sp$BLfamily!="Sulidae",]
respvar <- respvar[respvar$BLsciname %in% sp$BLsciname, ]

sp$BLsciname <- as.character(sp$BLsciname)
respvar$BLsciname <- as.character(respvar$BLsciname)
sp <- sp[, !names(sp) %in% c("Ncell","Nsite","Nrecord","Fyear","Lyear","Ncellcomb","BodyMassTamas")]

# number of cells per species
Ncell <- data.frame(BLsciname=names(table(respvar$BLsciname)), Ncell=as.numeric(table(respvar$BLsciname)))
sp <- merge(sp, Ncell, by="BLsciname")

# mean NA proportions per species
meanNA <- ddply(respvar, .(BLsciname), summarise, NAp=mean(mnNAprop))
sp <- merge(sp, meanNA, by="BLsciname")

# sort with PHYlabel
sp <- sp[order(sp$PHYlabel),]

## upload phylogeny
phylo <- read.nexus("Tree.213species.nex")

## data for bugs
Ntree <- length(phylo)
Ncell <- sp$Ncell
Nsp <- nrow(sp)       # 213 species in total

# species-level traits
LAT <- sp$absmidlat - mean(sp$absmidlat) # range mid latitude (centred across species)
LATrange <- sp$absmaxlat - sp$absminlat  # latitude range
MG <- ifelse(sp$mig.status%in%c("nomadic","not a migrant"), 0, 1) # 0=residents, 1=migrants
BM <- log10(sp$BodyMass)
NAperc <- 100*sp$NAp  # % NA

# calculate a covariance matrix from the tree (scaled to a height of 1)
varcov <- lapply(phylo, function(x){vcv(x, corr=T)})
# order by species name
varcov.order <- lapply(varcov, function(x){x[order(rownames(x)), order(colnames(x))]})

# save the inverse of the covariance matrix as an array
varcov.array <- array(NA, dim=c(Nsp,Nsp,Ntree))
invA <- array(NA, dim=c(Nsp,Nsp,Ntree))
for(k in 1:Ntree){
  varcov.array[,,k] <- varcov.order[[k]]
  invA[,,k] <- solve(varcov.array[,,k])
}

# check the order of the tree and data
ifelse(sp$PHYlabel==rownames(varcov.order[[1]]), "OK", sp$PHYlabel)
ifelse(sp$PHYlabel==colnames(varcov.order[[1]]), "OK", sp$PHYlabel)


## grid-level data on response and explanatory variables
resp.var <- matrix(NA, nrow=Nsp, ncol=max(Ncell))
# number of coefficients
Ncoefbtw <- 6
Ncoefwth <- 3
expv <- array(NA, dim=c(Nsp, max(Ncell), Ncoefwth-1)) # species by cells by predictors

## define spatial neighbourhood function
spneigh <- function(x,y,xrange=1,yrange=1){
  num<-length(x)
  cell<-data.frame(x=x,y=y,a=1:num)
  nearest<-nnear<-c(NULL)
  
  for(i in 1:num){
    subcell<-subset(cell,(cell$x<=cell[i,]$x+xrange)&
                      (cell$x>=cell[i,]$x-xrange)&
                      (cell$y<=cell[i,]$y+yrange)&
                      (cell$y>=cell[i,]$y-yrange)&
                      (cell$a!=cell[i,]$a))
    nearest<-append(nearest,subcell$a)
    nnear<-append(nnear,nrow(subcell))
  }
  return(list(Adj=nearest,Num=nnear))
}

## upload data for each species
for(i in 1:Nsp){
  
  sresp <- respvar[respvar$BLsciname==sp$BLsciname[i],]
  colnames(sresp)[3:4] <- c("Longitude","Latitude")
  
  # store response variable
  resp.var[i, 1:nrow(sresp)] <- sresp[, "mnTcoef"] # use mnTcoef for temp coefficients and mnPcoef for prec coefficients
  
  # store explanatory variables
  expv[i, 1:nrow(sresp), 1] <- abs(sresp[,"Latitude"]) - mean(sp$absmidlat) # grid cell latitude (centred with across-species mean mid lat)
  expv[i, 1:nrow(sresp), 2] <- 100*sresp[,"mnNAprop"] # mean % NAs in each cell
  
  # create adj and num for car.norm 
  x <- sresp$Longitude; y <- sresp$Latitude; a <- 1:nrow(sresp)

  result<-spneigh(x,y, xrange=1.5, yrange=1.5)
  if(length(result[[1]])<1){ result<-spneigh(x,y, xrange=2, yrange=2)}
  if(length(result[[1]])<1){ result<-spneigh(x,y, xrange=2.5, yrange=2.5)}
  if(length(result[[1]])<1){ result<-spneigh(x,y, xrange=3.5, yrange=3.5)}
  if(length(result[[1]])<1){ result<-spneigh(x,y, xrange=4.5, yrange=4.5)}
  if(length(result[[1]])<1){ result<-spneigh(x,y, xrange=5.5, yrange=5.5)}
  Adj<-result[[1]]
  Num<-result[[2]]
  Weights<-rep(1,length(Adj))
  (if(i==1){
    list.adj <- list(Adj); list.num <- list(Num); list.weights <- list(Weights)
  }
    else{
      list.adj <- c(list.adj, list(Adj))
      list.num <- c(list.num, list(Num))
      list.weights <- c(list.weights, list(Weights))
    })
}

l.adj <- sapply(list.adj,length)
l.num <- sapply(list.num,length)
l.weights <- sapply(list.weights,length)

mat.adj <- matrix(NA, nrow=Nsp, ncol=max(l.adj))
mat.num  <- matrix(NA, nrow=Nsp, ncol=max(l.num))
mat.weights <- matrix(NA, nrow=Nsp, ncol=max(l.weights))
for(i in 1:Nsp){
  mat.adj[i,1:l.adj[i]] <- list.adj[[i]]
  mat.num[i,1:l.num[i]] <- list.num[[i]]
  mat.weights[i,1:l.weights[i]] <- list.weights[[i]]
}


## define BUGS model ---------------------
cat("model
    {

    ### priors ###

    ## overall error
    tau ~ dgamma(0.01, 0.01)
    sd <- sqrt(1/tau)
    
    ## phylogenetic terms
    tauphy ~ dgamma(0.01, 0.01)
    sdphy <- sqrt(1/tauphy)
    lambda ~ dunif(0, 1)
    
    # tree sampling and lambda computation
    for(i in 1:Ntree){
     p[i] <- 1/Ntree
    }
    K ~ dcat(p[])   
    
    for(i in 1:Nsp){
      for(j in 1:Nsp){
        LAMBDA[i,j] <- 1 + (lambda-1)*(1-equals(i,j))
        TAU[i,j] <- tauphy*LAMBDA[i,j]*invA[i,j,K]
      }
    }
    
    for(i in 1:Nsp){
      mophy[i] <- 0
    }
    delta[1:Nsp] ~ dmnorm(mophy[], TAU[,])
    
    ## among-species coefficients
    # global intercept
    alpha ~ dnorm(0, 0.01)

    # for covariates
    for(i in 1:Ncoefbtw){
      b.btw[i] ~ dnorm(0, 0.01)
    }
    
    ## species specific terms
    for(i in 1:Nsp){
      # error terms
      tau.sp[i] ~ dgamma(0.01, 0.01)
      sd.sp[i] <- sqrt(1/tau.sp[i])
      
      # spatial terms
      tau.car[i] ~ dgamma(0.01, 0.01)
      sd.car[i] <- sqrt(1/tau.car[i])
      gamma[i, 1:Ncell[i]] ~ car.normal(mat.adj[i,1:l.adj[i]], mat.weights[i,1:l.weights[i]], mat.num[i,1:l.num[i]], tau.car[i])      
    }
    
    ## within-species coefficients
    # hyper parameters
    for(i in 1:Ncoefwth){
      hb.wth[i] ~ dnorm(0, 0.01)
      tau.b[i] ~ dgamma(0.01, 0.01)
      sd.b[i] <- sqrt(1/tau.b[i])
    }
    # each species
    for(i in 1:Nsp){
      for(j in 1:Ncoefwth){
        b.wth[i,j] ~ dnorm(hb.wth[j], tau.b[j])
      }
    }

    ### observations ###
    
    # for each species
    for(i in 1:Nsp){

      # species effect including covariates
      mu[i] <- alpha                      # overall intercept
             + b.btw[1] * LAT[i]          # absolute latitude midpoint - linear 
             + b.btw[2] * pow(LAT[i], 2)  # absolute latitude midpoint - quadratic
             + b.btw[3] * LATrange[i]     # absolute latitude range
             + b.btw[4] * MG[i]			      # migratory status
             + b.btw[5] * BM[i]			      # body mass
             + b.btw[6] * NAperc[i]       # mean % NA
             + delta[i]                   # phylogenetic autocorrelation term
      
      species[i] ~ dnorm(mu[i], tau)I(-10,10) # species effect
    
      # for each cell within each species 
      for(j in 1:Ncell[i]){

        m.resp[i,j] <- species[i] 
                      + b.wth[i,1] * (expv[i,j,1] - LAT[i])/LATrange[i]	        # abs latitude - linear (centred around range mean and divided by lat range)
                      + b.wth[i,2] * pow((expv[i,j,1] - LAT[i])/LATrange[i], 2) # abs latitude - quadratic
                      + b.wth[i,3] * expv[i,j,2]                                # % NA
                      + gamma[i,j]                                              # spatial autocorrelation term
  
        resp.var[i,j] ~ dnorm(m.resp[i,j], tau.sp[i])

      }

    }

    }",
    file="Model1.txt")


### modelling ---------------------
data <- list("Ncell","Nsp","resp.var","mat.adj","mat.num","mat.weights","l.adj","l.num","l.weights",
             "invA","Ntree","LAT","LATrange","MG","BM","NAperc","expv","Ncoefbtw","Ncoefwth")
parameters <- c("lambda","b.btw","hb.wth","b.wth","species","sd","sdphy","alpha","delta","sd.sp","sd.car","sd.b")
inits <- function(){
  gamma.init <- matrix(NA, nrow=Nsp, ncol=max(Ncell))
  for(i in 1:Nsp){
    gamma.init[i,1:Ncell[i]] <- rep(0, Ncell[i]) # zero-sum constraint 
  }
  list(tau=rgamma(1, shape=10, scale=0.1),
       tauphy=rgamma(1, shape=10, scale=0.1),
       lambda=runif(1, min=0, max=1),
       delta=rnorm(Nsp, mean=0, sd=0.1),
       alpha=rnorm(1, mean=0, sd=0.1),
       b.btw=rnorm(Ncoefbtw, mean=0, sd=0.1),
       tau.sp=rgamma(Nsp, shape=10, scale=0.1),
       tau.car=rgamma(Nsp, shape=10, scale=0.1),
       gamma=gamma.init,
       hb.wth=rnorm(Ncoefwth, mean=0, sd=0.1),
       tau.b=rgamma(Ncoefwth, shape=10, scale=0.1),
       b.wth=matrix(rnorm(Nsp*Ncoefwth, mean=0, sd=0.1), nrow=Nsp, ncol=Ncoefwth),
       species=rnorm(Nsp, mean=0, sd=0.1)
  )
}

### Run BUGS -----------------------
start <- Sys.time()
res<-bugs(data,inits,parameters, model.file="Model1.txt", n.chains=3,
          debug=F, n.iter=30000, n.burnin=10000, n.thin=4, DIC=FALSE,
          working.directory = getwd())
print(Sys.time()-start)

# save summary results
pdf("summary.pdf")
plot(res)
dev.off()
write.table(res$summary,"summary.csv",sep=",")
write.table(res$sims.array,"sims.csv",sep=",")
```
